# Large-scale proteomic analysis of T. spiralis muscle-stage ESPs identifies a novel upstream motif for *in silico* prediction of secreted products

**DOI:** 10.1101/2022.08.23.504907

**Authors:** Bradley Nash, William F. Gregory, Rhiannon R. White, Anna Protasio, Steve P. Gygi, Murray E. Selkirk, Michael P. Weekes, Katerina Artavanis-Tsakonas

## Abstract

The *Trichinella* genus contains parasitic nematodes capable of infecting a wide range of hosts including mammals, birds and reptiles. Like other helminths, *T. spiralis* secretes a complex mixture of bioactive molecules capable of modulating its immediate surroundings and creating a hospitable environment for growth, survival and ultimately transmission. The constitution of these excretory-secretory products (ESPs) changes depending on the tissue niche and the specific stage of parasite development. Unique to *T. spiralis* is a true intracellular stage wherein larvae develop inside striated myotubes. Remarkably, the parasite larvae do not destroy the host cell but rather reprogram it to support their presence and growth. This transformation is largely mediated through stage-specific secretions released into the host cell cytoplasm. In this study, we apply state of the art proteomics and computational approaches to elucidate the composition and functions of muscle-stage *T. spiralis* ESPs. Moreover, we define a commonly-occurring, upstream motif that we believe is associated with the stichosome, the main secretory organ of this worm, and can thus be used to predict secreted proteins across experimentally less tractable *T. spiralis* life cycle stages.

**Author Summary:** *Trichinella spiralis* is the only helminth parasite with a true intracellular stage. Newborn larvae penetrate the intestinal wall of the host, enter the circulation and preferentially infect muscle cells. Remarkably, they do not destroy the host cell but rather initiate a series of modulatory events that transform it into a ‘nurse cell complex’, a collagenated cyst that can persist for years. Each stage of *T. spiralis* development is guided by host-targeted secretions released by the worm directly into its immediate environment, mediating events such as immunoregulation, cell cycle control and angiogenesis. As such, these worm effectors hold therapeutic potential for chronic and autoimmune diseases. The composition of excretory-secretory products (ESPs) changes according to what the worm needs to accomplish and what tissue niche it is occupying at the time, with many deriving from the stichosome, the worm’s dedicated secretory organ. In this study, we characterise ESPs of muscle-stage *T. spiralis* larvae using proteomic and bioinformatic approaches and we define a regulatory motif associated with stichosome-derived proteins.

## Introduction

*Trichinella spiralis* is an obligate parasitic nematode and the leading cause of Trichinellosis (also called Trichinosis). During infection with *T. spiralis*, the new-born larva (NBL) of the parasite invades a terminally differentiated myofiber and resides intracellularly. The growth and development of the larva coincides with the reprogramming of the host muscle cell into a nurse cell for long-term inhabitation. The invading NBL and the host muscle cells undergo significant change during the process of parasite niche remodelling. The host cell re-enters the cell cycle, loses muscular specialisation, and gains a thick collagen capsule. The NBL undergoes significant growth, its cuticle thickens, and it develops the secretory organ, the stichosome, that matures by day 7 post-invasion[1,2]. The proteins secreted from the stichosome are hypothesised to mediate the transformation of a terminally differentiated muscle cell into a nurse cell complex, capable of supporting *T. spiralis* development[3].

Prior to the sequencing of the *T. spiralis* genome, the identification of *T. spiralis* excretory/secretory products (ESPs) using mass spectrometry was initially achieved using 2-D Electrophoresis for individual spot-based protein identification. The muscle-stage larva (MSL) ESPs were found to be highly tyvelosylated, isoform rich, and increasing in complexity during developmental progression [4,5]. Due to the complex composition of the ESPs, it is expected that the ESPs of the early MSL (eMSL) differ from the ESPs of the mature MSL (mMSL). However, isolation of the eMSL is hindered by their fragility and sensitivity to acidified pepsin up to Day 11 post-invasion [1,6]. The collection of ESPs from stages other than mMSL still requires the development of experimental procedures to effectively overcome these limitations.

Protein identification was greatly improved by the 2011 release of the *T. spiralis* draft genome (PRJNA12603)[7]. The proteomic annotations combined 3,262 genes identified from expressed sequence tag screens on Adult, MSL and NBL stages [8] with *ab initio* prediction using the software FgenesH and EAnnot. This laid the groundwork for the *T. spiralis* proteome containing a predicted 15,808 genes at the time of publication [7] and a later expansion to 16,380 genes. This advancement allowed for interpretation of the protein functions of MSL ESPs, identifying proteases, nucleases, glucose metabolising enzymes, and heat shock proteins from a single screen of only immunogenic bands from 1D-Electrophoresis [9].

Previously our lab identified the E2 Ubiquitin-conjugating protein TsUBE2L3 (T01_4545) from the ESPs of the mMSL and demonstrated its localisation to the stichosome of the muscle stage larvae [10]. Subsequent to this work, a second draft genome (PRJNA257433) has been constructed alongside transcriptomic profiling of pepsin-released MSL of *T. spiralis* and 15 other *Trichinella* species. The proteome annotations represent a dramatic improvement, combining transcriptomic data with homology-directed gene predictions, which were used to train further prediction of non-homologous genes [11]. In this study we sought to define the major activities of the ESPs to predict the host cell processes targeted by *T. spiralis* during muscle stage infection. We identified enriched pathways and abundant protein superfamilies that likely play a critical role in regulating the worm’s unique intracellular niche. We also sought to find a way to identify secreted, host-targeted genes bioinformatically. To this end we identified the PouStich motif which we hypothesise to be a stichosome-specific regulatory sequence.

## Results and Discussion

### Collection and annotation of ESPs

Mature L1 larvae were released from rat infected muscle by acidified pepsin digestion and cultured *in vitro*. The ESPs secreted into the culture medium were collected for a limited duration of 24 hours to minimise the risk of obtaining intracellular contaminants from dying worms. Secretions were concentrated and subjected to LC/MS/MS analysis as described in White et al. (2016) [10]. We identified 398 ESPs from either of 2 biological replicates and 208 proteins from both replicates **(Supplementary Table 1)**. To understand how the worm modulates the host environment, we proceeded to analyse bioinformatically the proteomic data to predict function, activity, localisation, and regulation of these secreted proteins.

### KAAS KEGG analysis reveals enrichment of glycolysis pathway components

KEGG Ontology (KO) terms were assigned to ESPs using the KEGG Automatic Annotation Server (KAAS)[14]. This analysis assigned KO terms to 48% of the ESPs (190/398) and only 26% of the *T. spiralis* proteome (3661/14,258) (**Figure 1**). Hypergeometric enrichment was used to determine whether members were present in the ESPs to a greater extent than would be expected based on their presence in the proteome for each KEGG pathway, which would suggest targeting of host cell pathways by *T. spiralis* ESPs (**Figure 2**).

**Figure 1.**
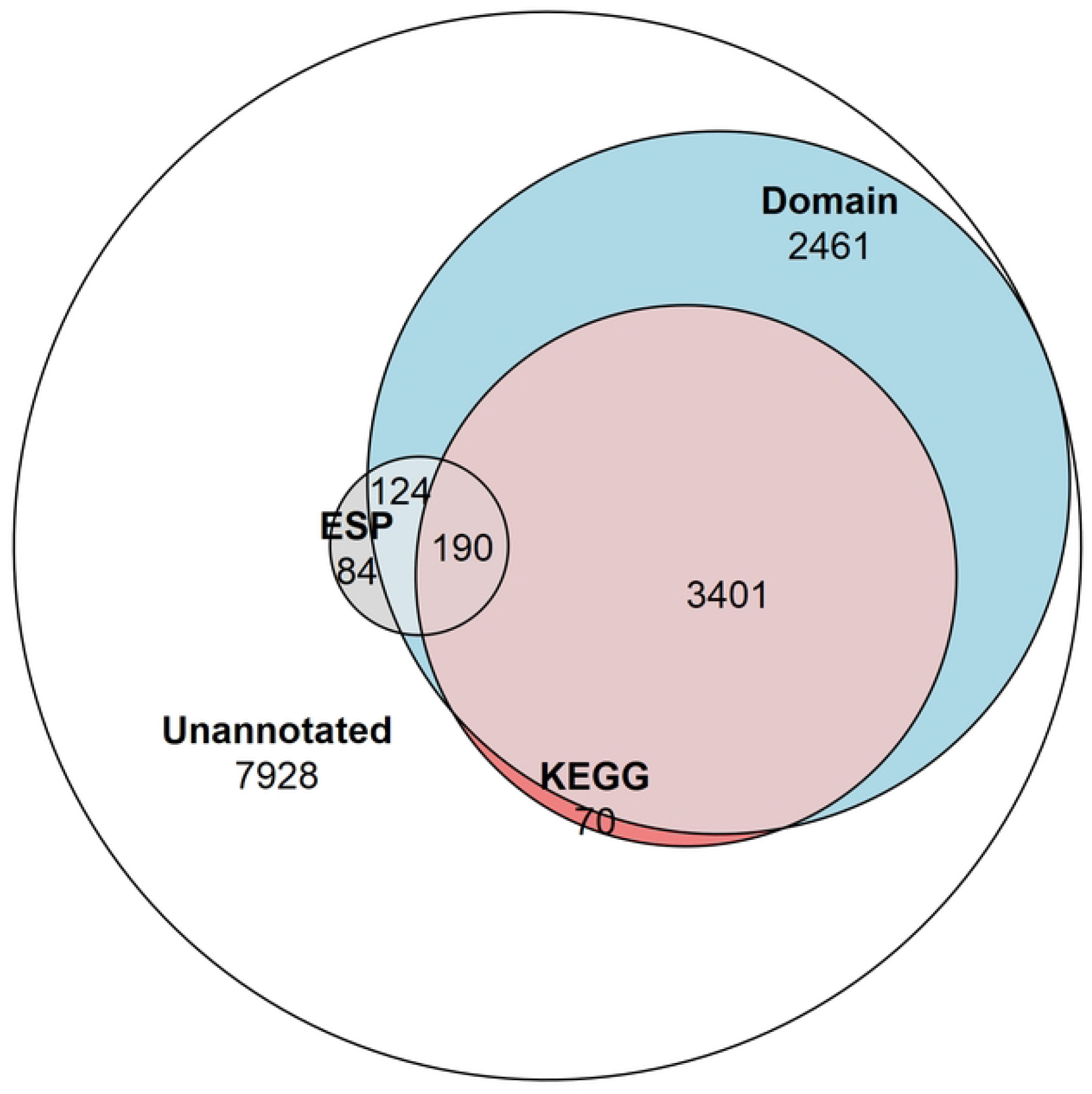
Venn diagram showing the proportion of the proteome and ESPs annotated. The number of NCBI Conserved Domain Search annotated proteins (6,176 total/314 ESPs), and KEGG Ontology annotated proteins (3,661 total/190 ESPs), and proteins not annotated by these approaches (7,928 total/84 ESPs).

**Figure 2.**
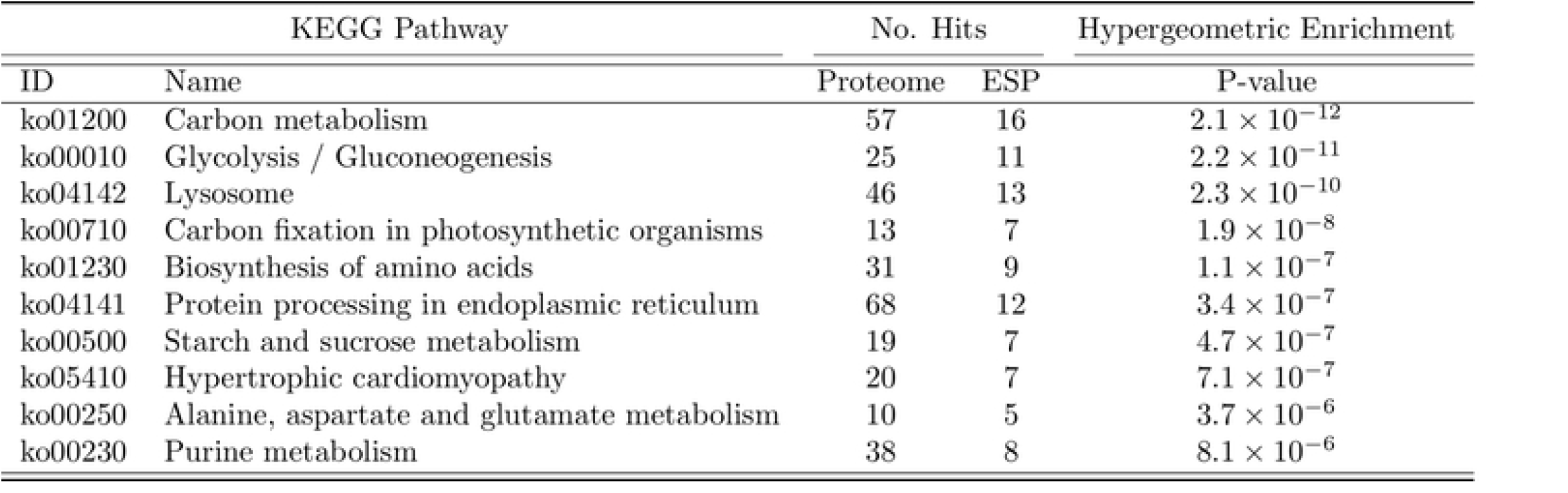
Top 10 enriched KEGG Pathways by excretory/secretory proteins (ESPs). Enrichment of KEGG Pathway members within the ESPs at a number of hits greater than expected based on the number of hits in the Proteome was determined using a hypergeometric test for enrichment (Phyper). The KEGG Pathways enriched by ESPs are ordered based on P value as an indication of the strength and reliability of the enrichment.

We observed enrichment of proteins involved in the KEGG pathway of hypertrophic cardiomyopathy, which was initially surprising. However, this condition stems from mitochondrial dysfunction [20], which is a hallmark of *T. spiralis*-infected muscle cells. ESPs were also enriched for carbohydrate metabolism pathways. Amylase, maltase and glucoamylase were identified in ESPs, which most likely serve a digestive function for larvae in the intestinal tract to convert starch and maltose to glucose, which could then be taken up by parasites. More interestingly, the entire glycolytic pathway was represented in ESPs with only 2 omissions, with entry into the pathway provided by glycogen phosphorylase (GP) rather than hexokinase (**Figure 3**). This suggests that *T. spiralis* mobilises glycogen stores, and as most animals store glycogen in skeletal muscle and the liver, this could be an important factor in growth and survival of parasites in muscle cells. Glycogen phosphorylase, phosphoglucomutase (PGM) and glucose-6-phosphate isomerase (GPI) are all represented in ESPs, which would drive conversion of glycogen to fructose-6-phosphate, However phosphofructokinase (PFK), a critical regulatory step for flux through glycolysis [21] was missing, either because it was below our level of detection, or suggesting that this step is performed by the host. Enzymes required for conversion of fructose-1,6-bisphosphate (F16BP) to phosphoenolpyruvate (PEP) were all present, with the notable exception of glyceraldehyde-3-phosphate dehydrogenase (GAPDH).

**Figure 3.**
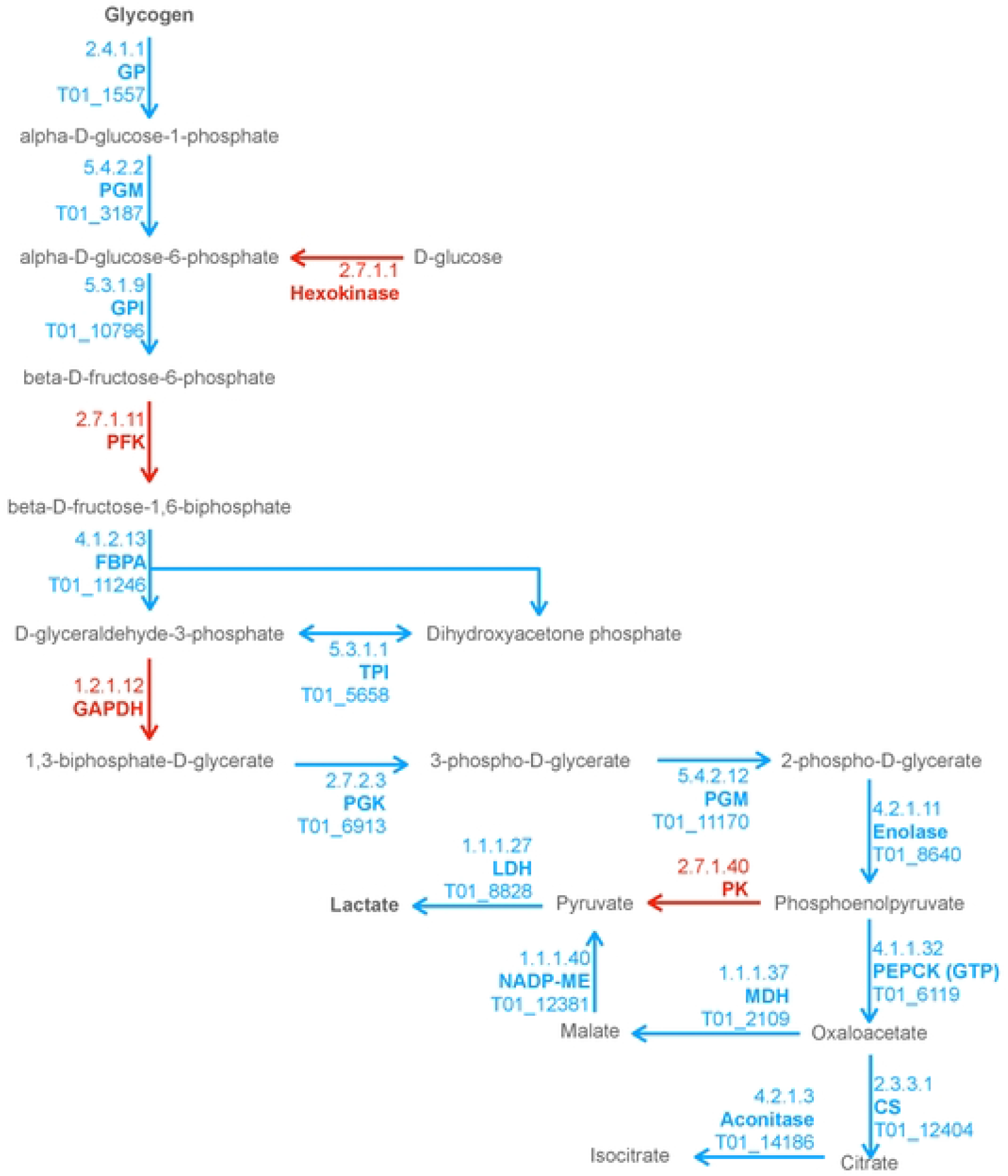
Combined excerpt of KEGG Pathways to illustrate ESPs that may be involved in catabolism of polysaccharide storages to lactate and TCA intermediates. The diagram illustrates the presence (blue with *T. spiralis* Gene_Stable_ID) and absence (red) of proteins (with KEGG Enzyme ID and Name) in the *T. spiralis* excretory/secretory proteins (ESPs). Metabolites are displayed in black, with arrows indicating anticipated flux direction. Proteins pertinent to the key stages Polysaccharide metabolism, Glycolysis, Malate dismutation and the TCA cycle have been indicated by differential background colours.

In anaerobic helminths, PEP can be converted to oxaloacetate by phosphoenolpyruvate carboxykinase (PEPCK) and then to pyruvate via malate by the process known as malate dismutation, rather than conversion of PEP to pyruvate by pyruvate kinase (PK) [22][23]. It was notable that all enzymes necessary for malate dismutation were present in ESPs, and that PK was missing. Enzymes that form the tricarboxylic acid (TCA) cycle were much less prevalent. Only 3 enzymes, malate dehydrogenase (MDH), citrate synthase (CS) and aconitase were found, which collectively could convert malate through to isocitrate. Further progression through the TCA cycle would therefore presumably be dependent on host enzymes.

It is possible that mobilisation of glycogen stores and secretion of glycolytic enzymes by *T. spiralis* alters host cell metabolism. Blood glucose levels are reduced from day 8 to 13 post infection, which can result in hypothermia [24,25]. Hypoglycaemia during the larval growth stage has been attributed to upregulation of insulin response genes, causing the glucose channel GLUT4 to be trafficked to the membrane, increasing glucose uptake. In this phase, nurse cells also build up glycogen stores [26], which may be necessary to sustain the exponential growth of the parasite [27].

Mature nurse cells exhibit a higher capacity for glucose uptake than L6 rat myoblasts during *in vitro* culture, which is dependent on both GLUT1/4 channels and H+/glucose import [28]. Nevertheless, we did not identify any *T. spiralis* secreted proteins which could mediate glucose uptake or drive glycogen synthesis. This suggests that both of these processes are regulated by host cells, whereas catabolism of glycogen may be driven largely by parasite secreted enzymes. Although some parasite glycolytic enzymes have been demonstrated to possess secondary functions [29][30], the near completeness of the glycolytic pathway represented by *T. spiralis* secreted proteins is strongly suggestive that they function primarily to regulate carbohydrate metabolism.

### Domain analysis reveals ESPs are particularly rich in proteases

Though beneficial for the confident identification of orthologs, KO terms can only be assigned to proteins that have a well-studied homolog, with a determined function, within a well-characterised pathway. These strict prerequisites resulted in less than half of the total ESPs being assigned a KEGG term by the KAAS. The NCBI Conserved Domain Search (NCDS) was used to expand the breadth of functional annotation to proteins containing domain signatures without requiring associated KEGG pathway involvement. The NCDS uses the Reverse-Position-Specific iterative Basic Local Alignment Search Tool (RPS-BLAST) approach to query a protein sequence against a collated database (Conserved Domain Database – v3.19 −58235 Position-Specific Scoring Matrices) of domains. The NCDS identifies domains which appear specific to sections of an input sequence, rather than the homology of a protein as a whole [12,31,32]. The application of the NCDS expanded the functional annotation of 190 (48%) ESPs to 314 (79%). All KEGG annotated ESPs also received domain annotation from the NCDS.

Multiple proteins were found to contain domains from the same superfamily, therefore the frequency of each domain superfamily was counted once per ESP presence and enrichment was scored to identify the ESP domains present at a greater frequency than expected from the proteome **(Figure 4)**. In concordance to the finding that carbon metabolism is a broadly enriched KEGG pathway, the AmyAc_family superfamily is identified as a highly enriched conserved domain. The AmyAc_family proteins are glycoside hydrolases whose substrates include the polysaccharides starch and glycogen which could act as an input for glycolysis.

**Figure 4.**
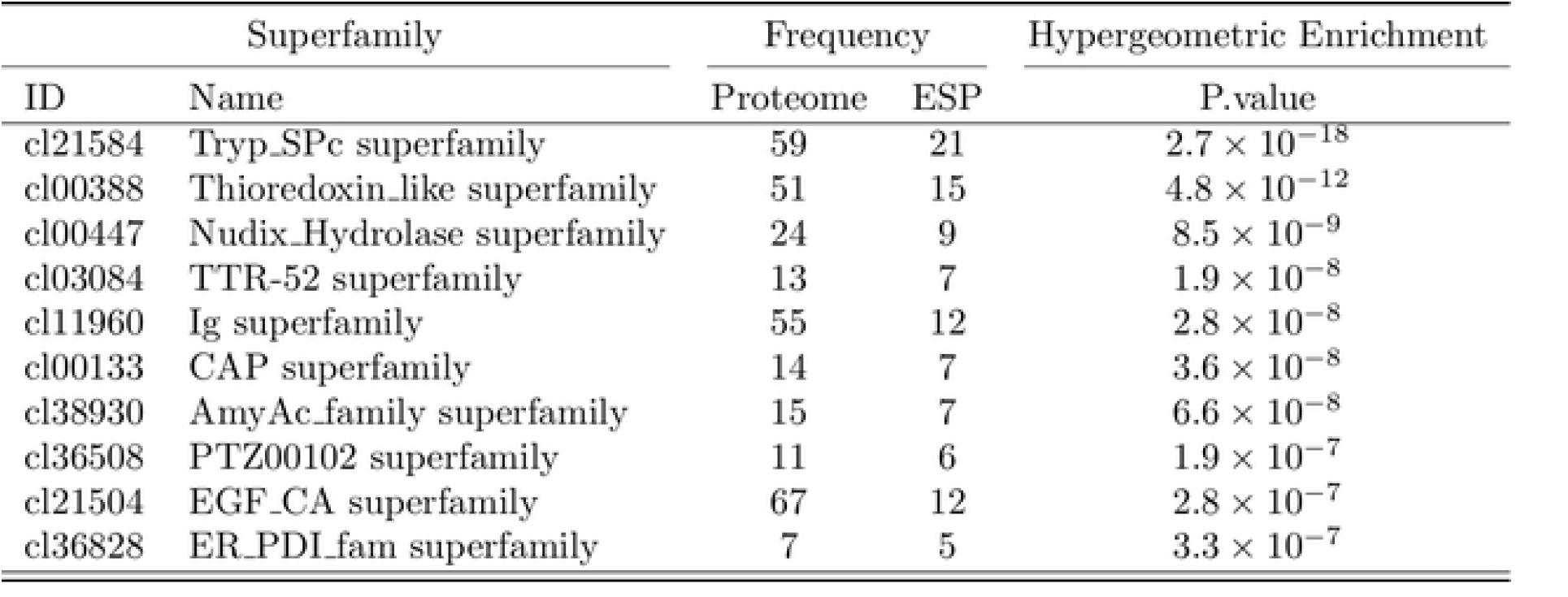
Top 10 most enriched domain superfamilies in the excretory/secretory proteins (ESPs). The occurrence of a domain associated with a superfamily term was detected by NCBI CDS and each occurrence of a superfamily domain was counted once per protein. The total incidence of proteins containing the domain within the excretory/secretory proteins has been summated and reported as Frequency. Superfamily names with their associated cluster ID are listed in order of P value, calculated using a hypergeometric test for enrichment (Phyper) to quantify the likelihood of the superfamily being enriched within the ESPs relative to their expected frequency based on the abundance of the superfamily within the proteome.

The proteolytic Tryp_SPc superfamily was the most prominent domain superfamily in ESPs. This family also contains the ESP with the most unique peptides, serine protease 30 (T01_15237). The activity of proteases within *T. spiralis* L1 larvae ESPs has been long established [33], with the proteases collected from MSL ESPs demonstrating different substrate specificity to the proteases collected from adult worm ESPs. Indicative of the collagen rich nurse cell complex, MSL ESPs more efficiently degrade gelatin and collagen, while adult ESP proteases are able to digest fibrinogen, haemoglobin and IgG [34]. The Tryp_SPc superfamily is the largest *T. spiralis* protease superfamily and is reported by the International Helminths Genome Consortium to have undergone a large family expansion, specifically within the parasites of Clade I (of which *Trichinella* is a member) [35]. We confirm the presence of reported MSL ESPs, including TS-15-1 (T01_13727) [36], TspSP-1 (T01_15237) [37], TsSP (T01_10159) [38], TsEla (T01_2909)[39] and TsChy (T01_6497) [40]. In a recent study, the Tryp_SPc domain from TS-15-1/T01_13727, found as an ESP, was able to elevate local collagen-I levels in mice after a 14-week treatment [36]. The study uncovered a mechanism dependent on protease activated cell surface receptor F2R-like trypsin receptor 1 (F2RL1) by which an ESP protease could alter host gene expression in construction of the nurse cell complex [41]. One could imagine that the collagen capsule surrounding the nurse cell is likely subject to frequent modification necessitating the function of enzymes involved in both its turnover and its incorporation. We noticed a striking level of diversity in the proteases present outside of the Tryp_SPc superfamily. We identified cysteine, serine and metalloproteases. We also identified an aspartate protease Ts-Asp/T01_3960 that has been detected in all life-stages except the NBL [42]. When not inhibited by pepstatin A, the protease preferentially interacts with the surface of intestinal epithelial cells rather than muscle cells [43]. To obtain MSL ESPs, the proteins were obtained by briefly culturing the MSL following release from the nurse cell by acidified pepsin. As a consequence, the ESPs collected during this time may constitute a mixture of proteins produced to function in the nurse cell and proteins produced in preparation for the next life-stage of intestinal colonisation. This finding would explain the presence of both proteases that have functions demonstrated to be relevant to both nurse cell and intestinal infection stages.

### ESPs are enriched in NUDT9-like nudix hydrolases

The nudix enzyme family encompasses over 80,000 proteins that hydrolyse a wide variety of substrates containing a nucleoside diphosphate linked to some other moiety, X [44]. They contain a characteristic consensus motif, generally G-x(5)-E-x(5)-[UA]-x-R-E-x(2)-E-E-x-G-(where U is a bulky hydrophobic residue). Members of the nudix family commonly have a wide range of physiological substrates ranging from (d)NTPs, oxidized (d)NTPs, non-nucleoside polyphosphates, and capped mRNAs, however the NUDT9 subfamily has only one known substrate, ADP-ribose (ADPR), which it converts to adenosine 5’-monophosphate (AMP) and ribose 5’-phosphate [45]. Thus, the NUDT9 family is also referred to as ADP-ribose hydrolases.

We identified a large extended family of nine NUDT9-type nudix hydrolases in *T. spiralis* muscle stage ESPs and a further four by searching the *T. spiralis* predicted proteome. In humans NUDT9 homologous sequences are found only in two proteins, the NUDT9 protein itself and in a highly homologous domain forming part of the calcium-permeable melastatin-related transient receptor potential cation channel (TRPM2 NUDT9H). *T. spiralis* nudix from ESP ranged in length from 279-517 amino acids and represent a set of evolutionarily diverging proteins compared to the canonical mammalian NUDT9, with extensions to both N and C termini containing predicted transmembrane domains which may determine protein localisation. Although the diversity in both sequence and length across the family of NUDT9 proteins was extensive, a phylogenetic tree could be constructed using an alignment of NUDT9 between the homology blocks **(Figure 5A)**.

**Figure 5.**
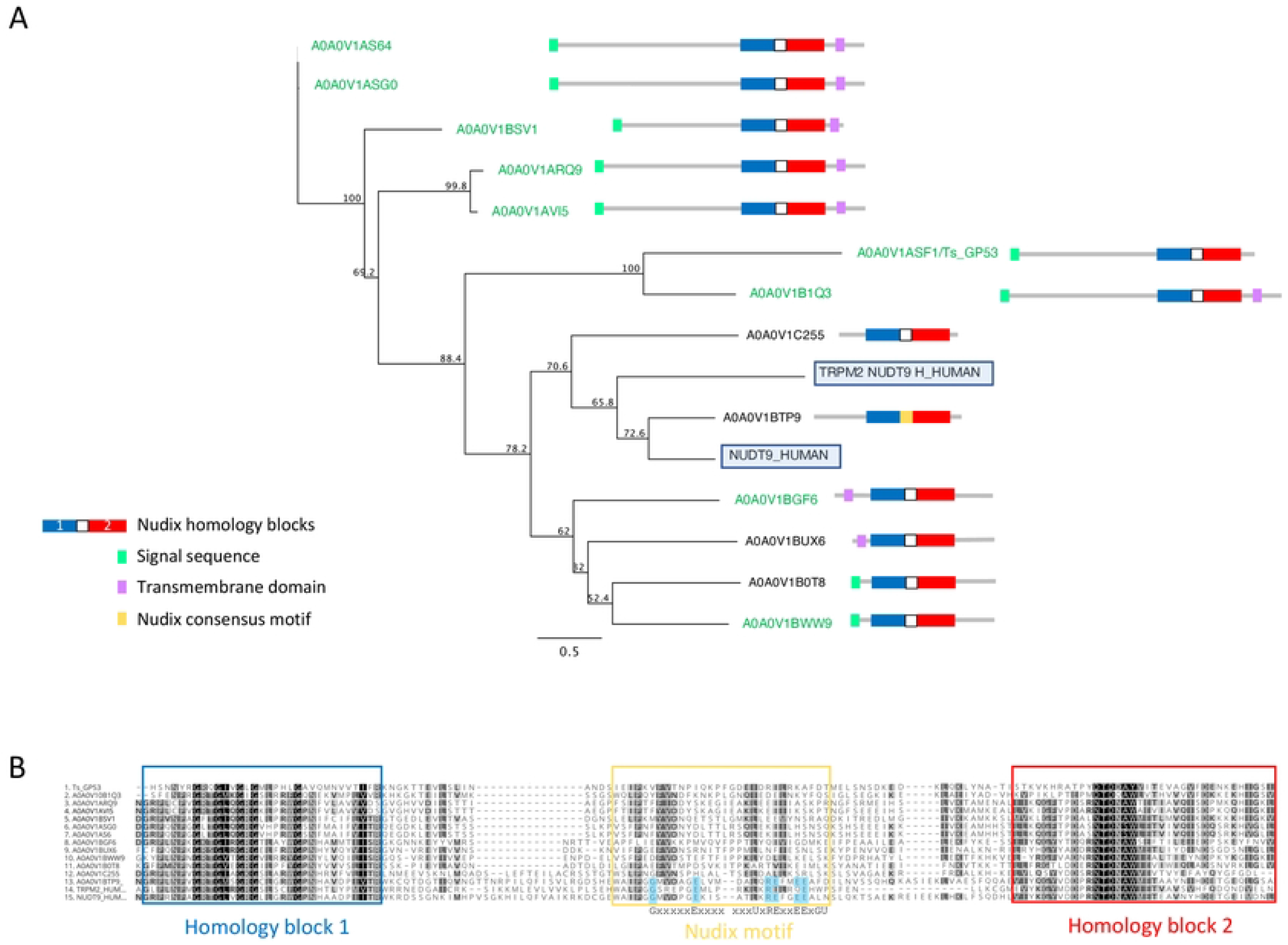
Alignment and phylogenetic analysis of *T. spiralis* NUDT9 homologs with their human counterparts. *A) Phylogenetic relationship of NUDT9 homologs. T. spiralis s*ecreted homologues are highlighted in green with human NUDT9 and TRPM2 NUDT9 homology domain boxed). The evolutionary tree was calculated using the maximum likelihood method implemented by the PhyML plugin for Geneious Prime. Proportions after 500 bootstraps are shown at each node. *B) Alignment of 13 T. spiralis NUDT9 homologues performed using ClustalW*. Residues conforming to the nudix motif are highlighted in blue. Conserved residues are coloured red. Residues with similar physiochemical properties are coloured yellow when >70% in each column.

**Figure 5B** shows an alignment of the *T. spiralis* nudix proteins with human NUDT9 and TRPM2 NUDT9H. Strong identity between these proteins and their mammalian counterparts is confined to two homology blocks (boxed in blue and red) either side of the nudix motif and thought to be involved in substrate binding [46]. A striking feature of the *T. spiralis* proteins is that all but one (A0A0V1BTP9) lack residues conserved within the consensus motif known to be essential for catalytic activity [47]. It is therefore likely that the *T. spiralis* proteins are catalytically inactive but still able to bind ADPR.

Human TRPM2 NUDT9H itself diverges from the nudix consensus motif and is unable to hydrolyse ADPR[48], however it retains the ability to bind ADPR, gating the channel which leads to Ca2+ influx into the cell [48]. The TRPM2 channel is activated under conditions of cellular stress as ADPR levels rise as a result of ADPR polymerization and degradation [49]. This leads to the release of inflammatory cytokines [50,51] and cell death if uncontrolled. This raises the possibility that *T. spiralis* nudix proteins bind free ADPR reducing or preventing Ca2+ influx via TRPM2.

It is also possible that *T. spiralis* nudix modulation of available ADPR may have broader effects within the cell such as a reduction in substrate availability for poly-ADPR polymerases (PARP). PARP conjugates ADPR to proteins playing important roles in gene transcription, chromatin organisation and stress response [52]. PARP-deficient mice are resistant to acute sepsis [53] and PARP inhibition prevents addition of mono and poly ADPR to NFkB, an important transcription factor regulating proinflammatory genes [54]. PARP also regulates other transcription factors such as AP-1, involved in the transcriptional response to oxidative stress[55]. Additionally, PARP has been shown to be involved in the promotion of the M1 macrophage phenotype associated with pro-inflammatory cytokines along with inhibition of M2 macrophages. [56].

These effects have several features in common with those reported for one of the most abundant secreted *T. spiralis* NUDT9 proteins A0A0V1ASF1 (also known as GP53/Ts53). It has been shown to suppress inflammatory responses in mouse models of sepsis [57] and colitis [58] and prevent death in mice from systemic administration of LPS [59].

Administration of recombinant GP53 results in the activation of M2 macrophages, perhaps via the modulation of PARP activity by GP53. It is therefore possible that *T. spiralis* nudix proteins regulate PARP activity by modulating ADPR availability interfering with the addition of ADPR to proteins and the synthesis of poly-ADPR.

### ESPs contain components of the extracellular matrix and enzymes associated with ECM assembly

After invasion of a muscle cell *T. spiralis* manipulates its environment outside the cell culminating in the formation of the collagenous capsule of the nurse cell which persists for the life-span of the host. Amongst the ESP proteins we find evidence that this manipulation continues after the nurse cell is formed. Of note is the secretion of two members of the Quiescin sulfhydryl oxidase (QSOX) family, proteins that catalyse disulphide bond formation. They are secreted into the extracellular environment by fibroblasts and are essential for the incorporation of laminin into the extracellular matrix (ECM) [60]. Importantly, we also see secretion of laminin by *T. spiralis*.

In addition, we see *T. spiralis* secreting a wide range of proteins that make up, or are associated with the ECM including six collagen family members, teneurin, and a homologue of the *C. elegans* protein, MUP-4.Teneurin is involved in the assembly of basement membrane collagen in *C*.*elegans*, interacts with dystroglycan, integrin and laminin and, when disrupted, leads to detachment of epidermis from muscle cells [61]. MUP-4 is essential for correct embryonic epithelial morphogenesis and maintenance of muscle position. It is also required for attachment of the apical epithelial surface and the cuticular matrix [62]. Another homologue of a cytoskeletal component, vinculin, is found within cell-cell and cell-ECM junctions and released by *T. spiralis*. We therefore hypothesise that ESP could contribute to the maintenance of the nurse cell within the muscle tissue niche during its long-term and stable infection.

### Abundance of ESPs predicted to localise to the host-parasite interface

ESPs are released directly into the host cell and have been demonstrated to localise to different organelles of the nurse cell [63]. To complement functional annotation of the ESPs we also used WoLF PSORT to predict subcellular localisation of ESPs within the infected host cell based on inherent protein sequence properties. WoLF PSORT assigns localisation of a protein by similarity to the features of the nearest neighbour class with a known subcellular localisation [16]. WoLF PSORT predicted plasma membrane (32%) and extracellular (25%) localisation based on protein features for greater than half the identified ESPs. The analysis indicates that a significant proportion of the ESP is expected to be trafficked from *T. spiralis* through the nurse cell, to presumably mediate interaction with the host and the nurse cell complex **(Figure 6)**. The importance of the ESPs that function at the nurse cell complex interface is supported by enrichment of classically extracellularly functioning conserved domains. We identify enrichment in the ESPs for the Ig superfamily and EGF_CA superfamilies that typically occur on the cell surface. The finding supports that despite the parasite’s intracellular confinement, many ESPs may be trafficked to the plasma membrane and contribute to the extracellular environment.

**Figure 6.**
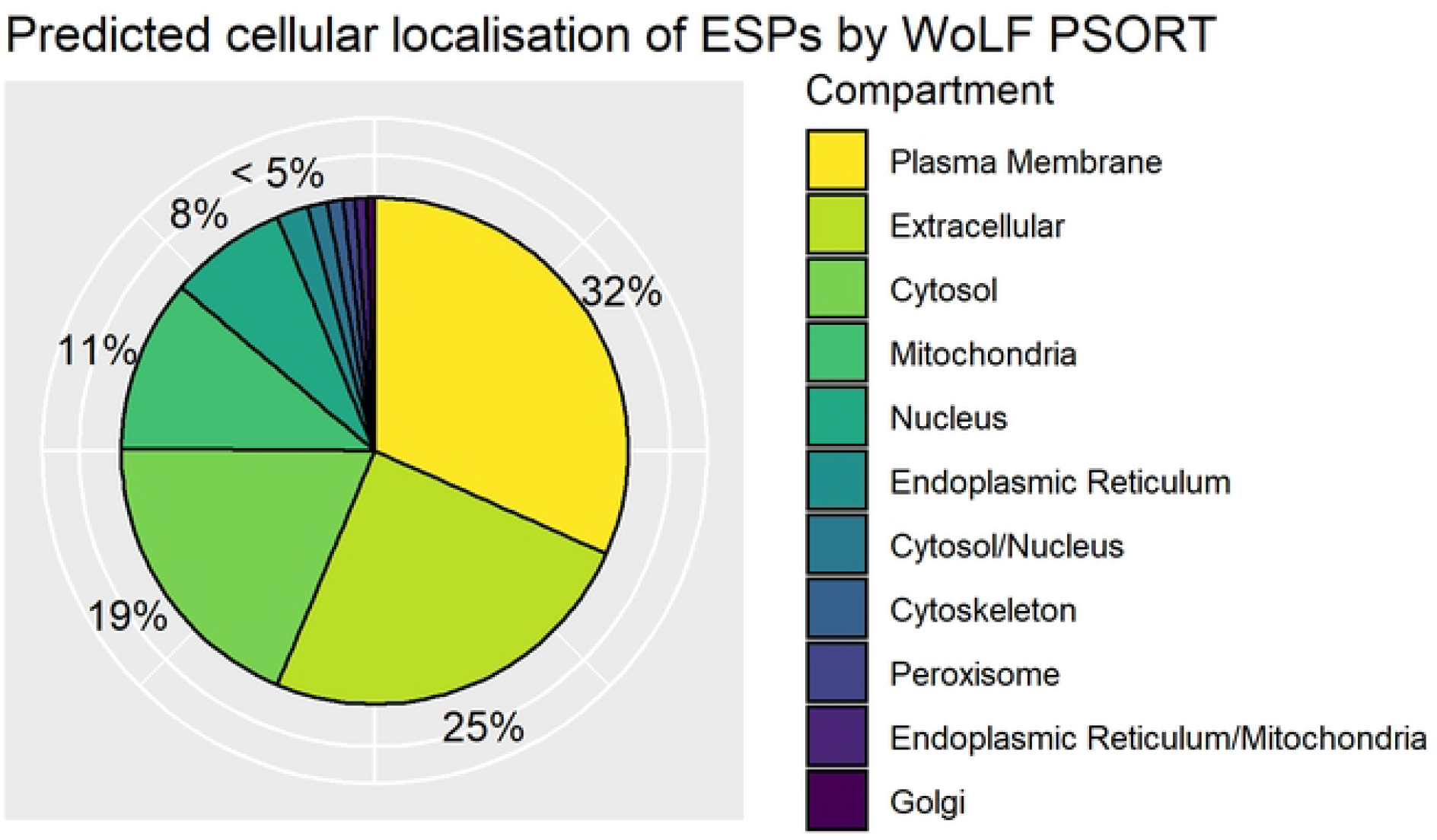
Distribution of WoLF PSORT predicted excretory/secretory (ES) protein localisations. Pie chart of top predicted subcellular localisations for ES proteins by WoLF PSORT as a percentage of the ESPs. Dual localisations indicated by “/” represent proteins expected to shuttle between two compartments.

### Only a minority of ESPs contain classical secretion signal peptides

Since our identified ESPs are secreted, one would expect them to contain signal peptides. Surprisingly, we found that only 36% (143 proteins) contained a recognisable N-terminal secretion signal using SignalP6.0 [15]. The low incidence of secretion signals hints at the complexity of the ESPs collected from mMSL culture supernatant and suggests they have classical secretion dependent and independent origins. Our results may under-represent the number of ESPs containing secretion signals, since their recognition is entirely dependent on the N-terminal sequence of the protein. Many of the proteomes’ gene models were constructed from transcriptomics data and protein sequences have not been confirmed experimentally. Therefore, up or down-stream mis-predictions of translation start sites may hinder the detection of secretion signals.

We found that the majority of proteins anticipated to localise extracellularly also contain a canonical secretion signal, again detected by SignalP6.0 (**Supplementary Table 1)**. However, 25% of predicted extracellular proteins and 66% of predicted plasma membrane localising proteins did not. Despite all the ESPs being collected externally from the nematode, the variable requirement for a secretion signal is still poorly understood. In *T. spiralis*, the stichosome is a dedicated secretory organ composed of α and β-stichocytes that contain respective α and β-granules. ES antigens are detected within the granules of the stichosome and predominantly exit from the mouth of the organism [64]. However, non-canonically secreted proteins may also be transported to the host cell as components of extracellular vesicles (EVs) that have been widely shown to have immunomodulatory effects such as upregulating anti-inflammatory cytokines (IL-10, IL-4, IL-13 and TGF-B) and Th2-associated transcription factors Foxp3 and GATA3 [65]. Given their immunomodulatory effects, MSL EVs must be capable of transporting proteins from the worm to the cell surface where they can interface with the immune system. A recent study that proteomically profiled EVs derived from MSL-stage *T. spiralis* identified 753 proteins [66]. Of these, 175 were also present in our dataset (**Supplementary Table 1)**. The overlap is perhaps not surprising given the fact that our ESP preparation would be expected to capture EVs. In the interest of understanding *Trichinella* biology, future studies to determine the subset of ESPs that are derived from the stichocytes and the subset of ESPs that originate from EVs of the cuticle would be greatly beneficial.

### Identification of a novel ESP-associated motif

The development of the dedicated secretory organ of *T. spiralis* does not progress until after the NBL establishes itself in the host muscle cell suggestive of an important role in nurse cell development. Since a large proportion of the ESPs are expected to originate from the stichosome, we hypothesised that the ESPs could be used to identify a stichosome-specific regulatory motif. A regulatory motif enriched upstream of genes identified as ESPs could, in turn, be used to predict stichosome-derived, secreted proteins across the life-cycle. Since ESPs are impossible to collect in sufficient quantities and properly study during the earliest stages of MSL development, such a predictive tool would be valuable in understanding how *T. spiralis* initiates muscle reprogramming.

We searched for a motif enriched in the region of 1000 bp upstream from the Transcription Start Site (TSS) of the 398 identified ESPs. The equivalent 1000 bp region was recovered for each of the 14,258 genes in the PRJNA257433 *Trichinella spiralis* genome [11] and used as background control sequences. The Homer2 software [19] was used to construct Hidden Markov Models (HMMs) for motifs that appear more frequently in the “regulated” target dataset (ESPs) compared to the background of “unregulated” sequences (Reference Proteome). Motif-direction outside of the core promoter region does not significantly influence the transcription regulating effect of a transcription factor on a gene [67], hence the motif search was performed on both strands.

The Homer2 motif search using known transcription factor binding sites, including those from the nematode model organism *C. elegans*, failed to identify any significantly enriched motifs. Given the absence of a stichosome in *C. elegans*, this result was not discouraging and highlights the divergence of regulatory processes in the genetically distinct nematode Clades. To ensure the exclusion of false positive hits, we interrogated ESPs with 2 or more unique peptides. Only 1 *de novo* motif was significantly enriched upstream of the TSS of these ESP genes. This motif has the highest similarity to the known human POU Class 2 Homeobox 3 (HsPou2f3) binding site (MA0627.2/Jaspar(0.761) and is highly enriched in what we hypothesise to be stichosome-derived proteins, so we named it PouStich (**Figure 7**).

**Figure 7.**
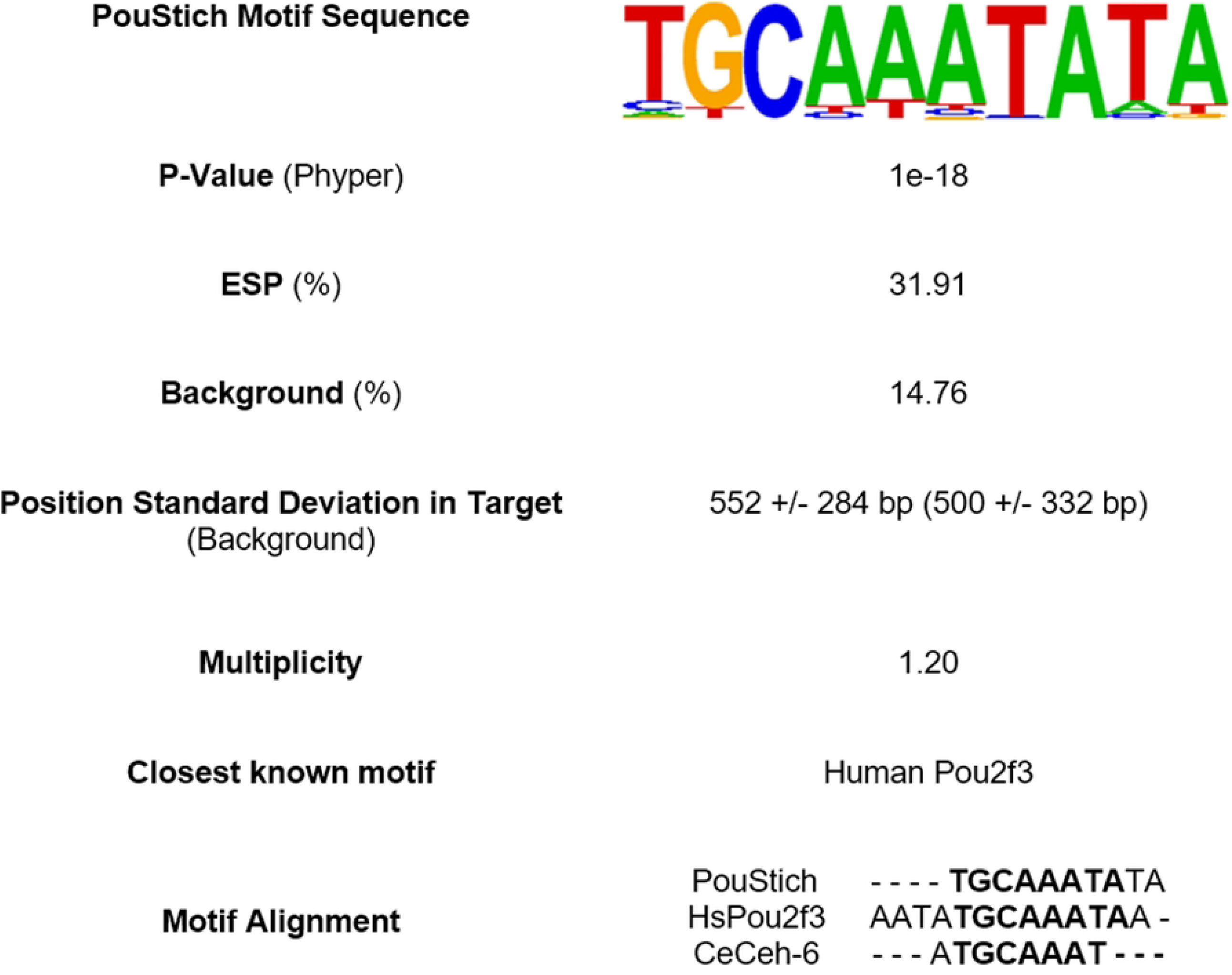
Summary of top Homer2 *de novo* motif search hit enriched upstream of excretory/secretory proteins (ESPs). The PouStich motif sequence is displayed as a Hidden Markov model logo. Beneath is the percentage of PouStich motif occurrence in the 1000bp upstream of the transcription start site (TSS) of ESPs compared to the “Background” of all *T. spiralis* protein coding genes and the P-value associated with the enrichment. The position of motif occurrence in distance from the TSS in both sample populations is indicated with +/-Standard deviation. The frequency of which the motif occurs multiple times in one upstream region is represented by the Multiplicity. The table also identifies the nearest known motif database hit and nearest *C. elegans* homolog motif as an alignment.

### Prediction of PouStich motif-binding proteins

BLASTp was used to identify proteins from *T. spiralis* that are similar in amino acid sequence to HsPou2f3 and therefore, likely to bind the PouStich motif. We identified Pou domain, class 3, transcription factor 2-A protein (TsPou3f2-a) as the closest *T. spiralis* hit to HsPou2f3. However, the human and the *T*.*spiralis* Pou domain proteins fall into different classes and appear evolutionarily distant. To explore a more similar background, BLASTp was used to interrogate the nematode *C. elegans* and identified the class 3 Pou domain containing transcription factor ceh-6 (CeCeh-6) (**Figure 8A and B**).

**Figure 8.**
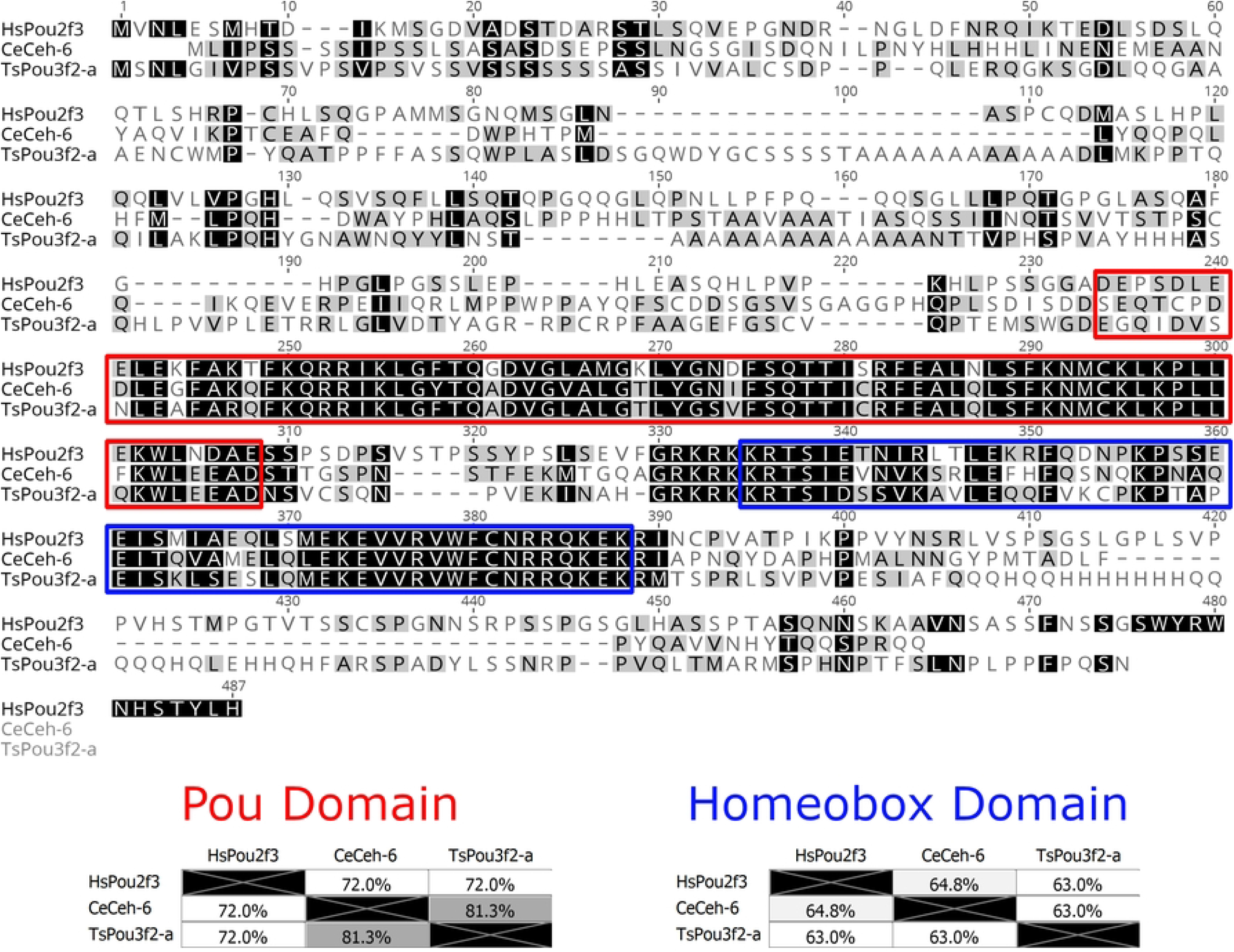
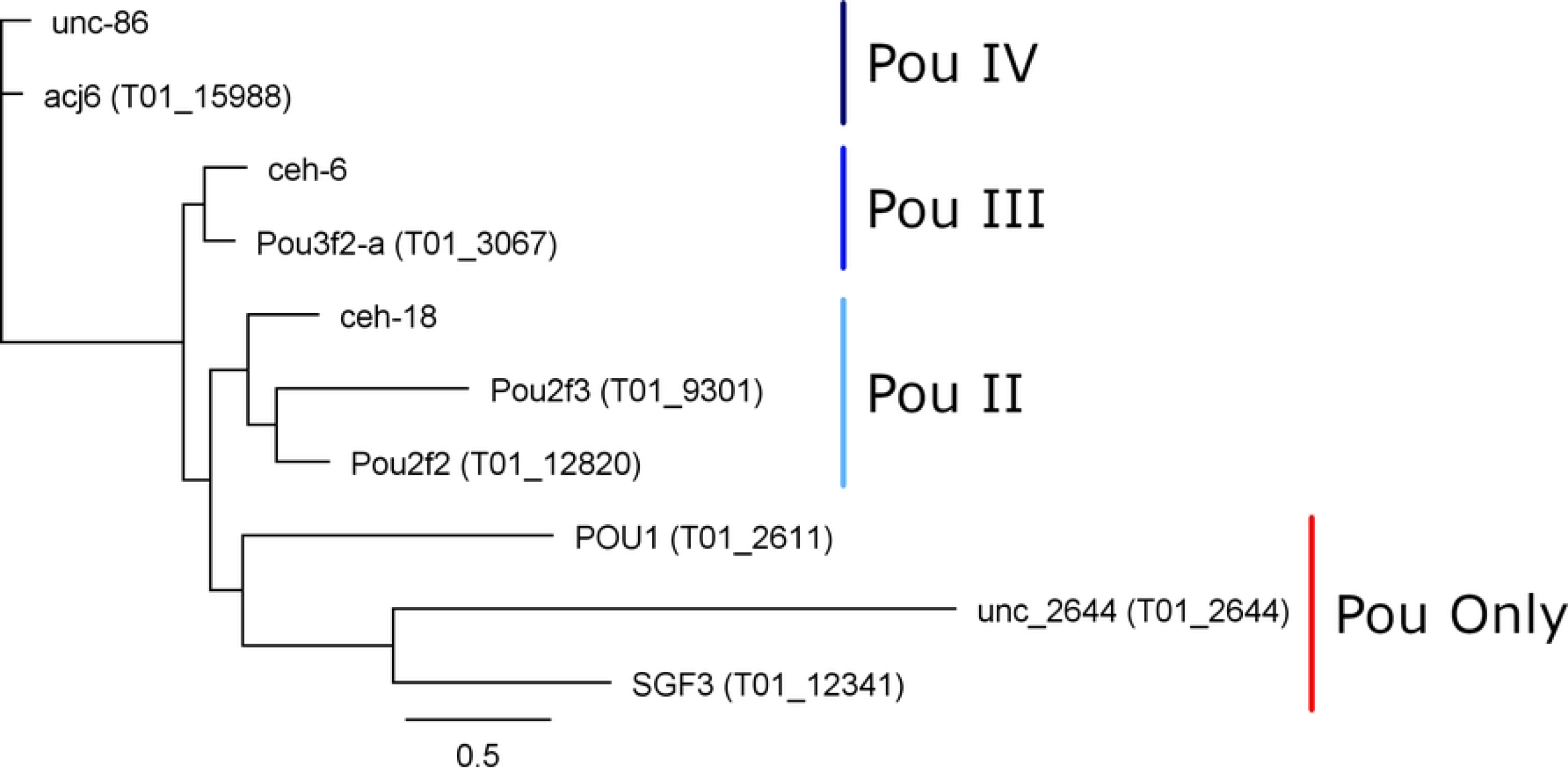
Pou domain containing octamer motif binding proteins. *A) MUSCLE alignment of the Human Pou2f3 with the closest C. elegans and T. spiralis BLASTp hits*. Amino acid shading highlights similarity. In the red box is the Pou domain and in the blue box is the homeobox domain, the two essential conserved regions of Pou-domain containing octamer motif binding proteins. *B) Phylogenetic tree from alignment of the T. spiralis and C. elegans Pou domains*. The tree was constructed with Phyml from aligned Pou domains across all sequences shown above using the Whelan and Goldman substitution model. The Pou domain type is annotated adjacent to the tree groupings. *T. spiralis* genes are included in brackets next to the gene name, while only *C. elegans* gene names are provided.

CeCeh-6 is the only class 3 Pou domain-containing protein identified in *C. elegans* and has a demonstrated role in proper excretory cell differentiation. Class 3 Pou domains are conserved across many phyla as an excretory/secretory cell regulator and are also implicated in the nervous system [68]. *C. elegans* has 3 Pou domain containing homeobox genes, with CeCeh-6 being the sole POU-III family member. Using a lacZ reporter, CeCeh-6 expression has been localised to discrete neural cell populations and the excretory cells. An effective null mutant for CeCeh-6 caused embryonic and larval lethality with larvae developing bodily vacuoles. The phenotype could be mimicked by laser ablation of the excretory cells, concluding that larval lethality was secondary to impaired excretory cell function [68]. The involvement of CeCeh-6 in excretory cell function in *C. elegans* is encouraging, as it supports the idea that the PouStich motif may mediate ESPs production.

Pou domain-containing proteins typically interact with DNA sequences through the combination of the Pou domain and the Homeobox (Hox) domain **(Figure 8A)**. The TsPou3f2-a sequence has 81% identity with CeCeh-6 across the Pou domain (red) but only 72% identity with HsPou2f3. Across the Hox domain (blue) TsPou3f2-a shares 63% identity with both HsPou2f3 and CeCeh-6. Taken together the results indicate that the Pou domain of TsPou3f2-a would bind a sequence more similar to the CeCeh-6 Pou interaction motif than that of HsPou2f3.

Using the NCDS, we identified a further 6 Pou domain-containing proteins from the *Trichinella spiralis* reference proteome and found that each *C. elegans* Pou domain containing protein has an identifiable homolog in *Trichinella spiralis*. This could suggest that nematodes require a minimum of three Pou domain-containing proteins of Class IV, Class III and Class II. In *Trichinella spiralis* however, a presumed duplication has resulted in the gain of a second Class II Pou domain containing protein. One Class II protein may function as a required nematode factor similarly to *C. elegans* CeCeh-18, while the other Class II protein might regulate the stichosome, an organ absent in *C. elegans*.

### Using the PouStich motif to predict novel ESPs

Hypothesising that the PouStich motif is predictive of stichosome origin, we used Homer2 to search for the PouStich motif in the region upstream of all reference proteome genes. We identified 2,157 genes containing the PouStich motif within 1000bp from the TSS as a direct match or with 1 tolerable mismatch (Motif Score > 9.216498), including 127 ESP genes from this study **(Supplementary Table 2)**. From the identified ESPs the two proteins with the highest unique peptide number, mcd-1 and serine protease 30, contained upstream PouStich motifs as did secreted from muscle stage larvae 3, sml-3 (T01_15139). The motif was also found upstream of proteins that were anticipated as ESPs but unconfirmed by peptides, such as sml-5 (T01_15780) and the glycolytic enzyme glyceraldehyde 3-phosphate dehydrogenase, gpd-1 (T01_7457). The proteins mcd-1, sml-3, and sml-5 have been demonstrated to localise to the MSL stichosome [69,70].

Not all genes expressed within the stichosome would be secreted as ESPs, some would likely be required for maintenance and differentiation of the secretory organ itself. It is likely that additional factors are required to identify the true ESPs amongst the motif containing genes. Of the 2,157 PouStich motif-regulated genes we detected co-occurrence of a Secretion Signal in only 175, 61 of these were also identified in our proteomic screen **(Figure 9 and Supplementary Table 1)**. It remains to be seen whether the other 114 proteins are secreted, supporting our hypothesis that the PouStich motif is an important regulator of the secretory proteins of *T. spiralis*.

**Figure 9.**
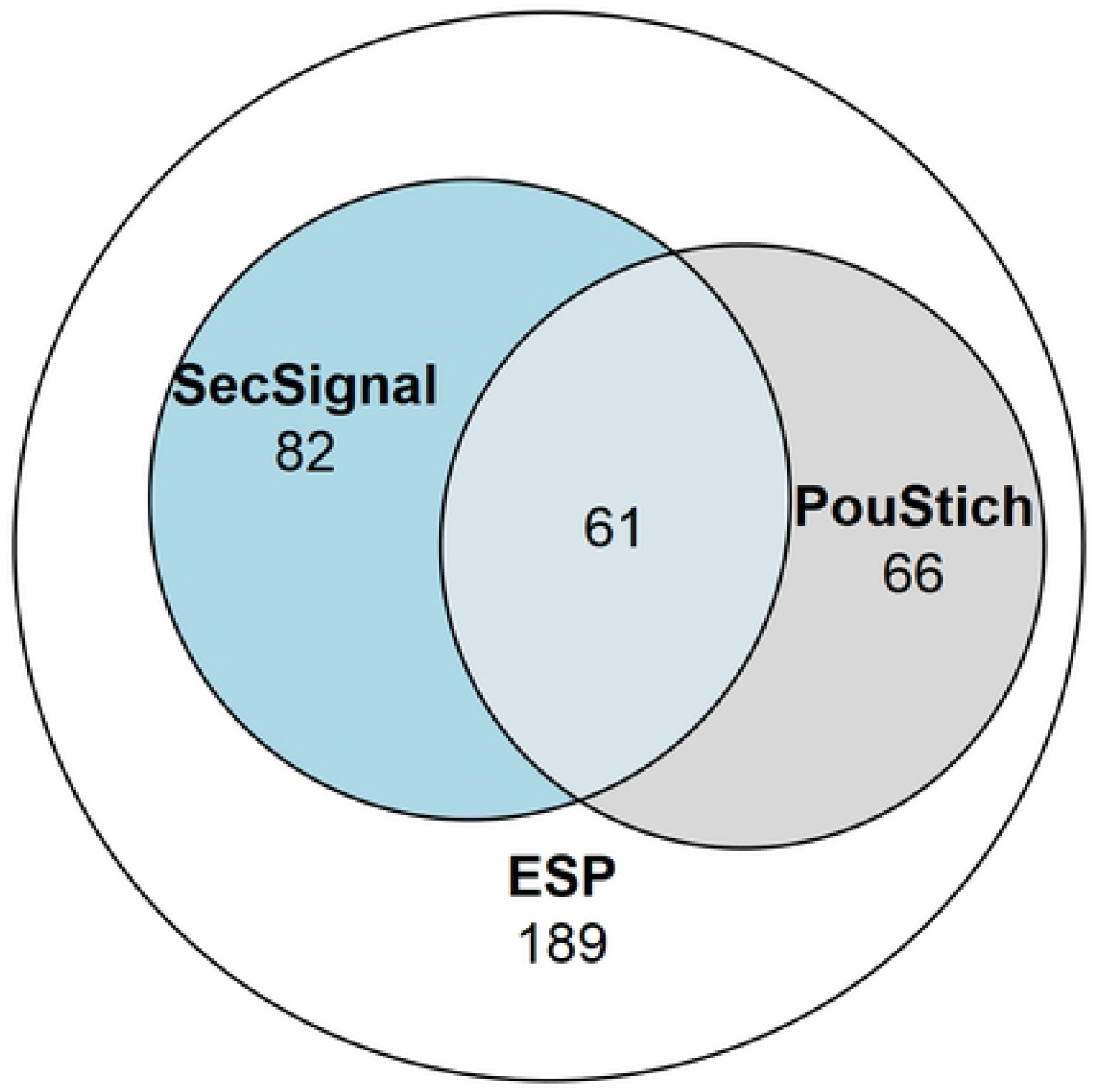
Venn diagram showing the proportion of the ESPs annotated with motifs. 398 ESPs were identified in this study, of which 143 contained a SignalP6.0 detected Secretion Signal, and 127 ESPs contained a Poutich motif upstream of the transcription start site. The 61 ESPs that contain a Secretion Signal and PouStich motif are a subset of high confidence predicted ESPs.

### Conclusions and Perspectives

Recent advancements in the construction of the *T. spiralis* proteome has enabled more extensive analysis of the secretome across the lifecycle. Combining these advances with modern protein annotation software allowed us to gain insight into the functions of the ESPs involved in this unique host-pathogen interface. We observed that the mMSL secretome is enriched for proteins predicted to affect metabolism. Furthermore, we identify superfamilies within the ESPs that predict multiple mechanisms through which the MSL can regulate the host immune system and remodel the extracellular environment during infection. We use the genomic context of ESPs identified in this study to highlight a novel motif, the PouStich motif, enriched upstream of their TSS. Though more experimental validation is required, the PouStich motif represents an exciting possibility that a portion of the ESPs are regulated by a common motif. Therefore, the PouStich motif in combination with other ESP associated patterns could allow the identification of ESPs from other currently inaccessible developmental stages.

## Materials and methods

### ESP preparation and LC/MS/MS analysis

ESP were collected and processed as described in White et al. (2016) [10]. Briefly, in two separate biological replicates, mature L1 larvae were released from rat infected muscle by acidified pepsin digestion and maintained in a culture flask in serum-free medium. The ESPs secreted into the culture medium were collected for 24 hours, concentrated using a 10 kDa protein filter, digested into peptides using trypsin, enriched and subjected to LC/MS/MS analysis. MS2 spectra were searched against a composite database derived from the UniProt *Trichinella spiralis* ‘one protein per gene’ reference proteome (14,258 genes, UP000054776), and potential contaminating proteins derived from pig trypsin, human and the Rattus Norwegicus UniProt proteome (download: 26/04/2022). Peptide spectral matches were filtered to a 2% false discovery rate using the target-decoy strategy combined with linear discriminant analysis. The mass spectrometry proteomics data will be deposited to the ProteomeXchange Consortium (http://www.proteomexchange.org/) via the PRIDE partner repository.

### Bioinformatic Analysis

Each identified ESP was queried against the Conserved Domain Database (v3.19 −58235 Position-Specific Scoring Matrices) using the NCBI Conserved Domain Search (https://www.ncbi.nlm.nih.gov/Structure/cdd/wrpsb.cgi?) with a significance threshold E-value of < 1×10^−2 [12]^. Gene ontology analysis was performed with gProfiler2 using the organism trspirprjna257433 [13]. Kyoto Encyclopedia of Genes and Genomes (KEGG) ontology (KO) terms were assigned to proteins using the KEGG Automatic Annotation Server (KAAS) (https://www.genome.jp/kaas-bin/kaas_main) using the Bi-directional Best Hit Assignment Method from Blast results against the default eukaryotic representative gene set with the additional inclusion of *T. spiralis* as a reference organism [14]. Canonical Eukaryotic secretion signals were detected at the N-termini of peptides using the SignalP-6.0 server (https://services.healthtech.dtu.dk/service.php?SignalP) [15]. WoLF PSORT

(https://wolfpsort.hgc.jp/) cellular localisations were determined by reporting the highest scoring localisation for each protein using the Animal organism type [16].

### Statistical analysis

The R function phyper was used to perform hypergeometric testing for enrichment. The P values were calculated using the script as follows: phyper(q - 1, m, n - m, k, lower.tail = FALSE) where q and m are constant and equal to No. Hits in the Subject and the Background respectively and while n and k are the total members of the Background and the Subject respectively. The arguments q – 1 and lower.tail = FALSE tests the hypothesis that enrichment is greater than or equal to the test population. Therefore, the P-value represents the probability that the hits within the subject population are greater than expected based on the hits in the background population as a result of chance, whereby P < 0.05 suggests enrichment.

### Sequence analysis

Sequence analysis was performed using Geneious 11.1.5. Sequences were aligned using Muscle 3.8.425. A phylogenetic tree was produced from aligned sequences using the PhyML 3.3.20180621 plugin with the Whelan and Goldman substitution model [17].

### Identification of enriched motifs

The 1000bp region upstream of the Transcription Start Site (TSS) was retrieved for each Reference proteome protein using BioMart (https://parasite.wormbase.org/biomart/martview/) hosted on WormBase Parasite [11,18]. For Homer2 (http://homer.ucsd.edu/homer) [19] input sequences, all the recovered sequences were used as the background, while the subject was the subset of sequences from identified ESP genes. The motif search was performed at a higher stringency level of proteins with 2 or more unique peptides to minimise the inclusion of false positive hits. The findMotifs.pl function was used to search for significantly enriched (P value < 1e-11) de novo motifs across all stringency levels, using -fasta to specify the custom background file and using fasta as the placeholder organism. Motif occurrence in the genome was determined using the -find option with the output ‘.motif’ file, to search the background sequences for the significantly enriched PouStich motif.

## Acknowledgements

This work was supported by a Wellcome Trust Career Development Award [WT085054MA], a BBSRC Project Grant [BB/R001642/1] both held by KAT and an MRC Studentship awarded to RW. BN was supported by a Department of Pathology PhD Studentship and Rosetrees Trust PhD Consumables [M703] and PhD Plus Grants [PhD2021/100037]. MPW is supported by Medical Research Council Project Grant MR/W025647/1 and SG is supported by an NIH RO1 grant [GM067945]. For the purpose of open access, the author has applied a CC BY public copyright licence to any Author Accepted Manuscript version arising from this submission.

